# Fire severity drives shifts in ectomycorrhizal fungal communities and Pseudotsuga menziesii seedling performance

**DOI:** 10.1101/2025.08.14.670368

**Authors:** Dustin D. Lower, Laura M. Bogar

## Abstract

**Background and Aims:** High-severity wildfires are increasing in western North America, disrupting soil conditions and mutualisms between plants and ectomycorrhizal fungi (EMF), which support seedling nutrient uptake, stress tolerance, and survival. This study investigates how fire severity and recency affect fungal communities and *Pseudotsuga menziesii* (Douglas-fir) seedling performance, and whether live-soil inoculation or stress priming improve post-fire outcomes.

**Methods:** In a greenhouse experiment, Douglas-fir seedlings were grown in soils from high-severity burns, recent and historic low-severity burns, and unburned sites in the Sierra Nevada. Each soil was applied as a live or autoclaved inoculum. Drought stress and abscisic acid priming treatments were imposed before a final 3-week dry-down. We measured seedling biomass, chlorophyll fluorescence (Fv/Fm), EMF colonization, and fungal community composition using ITS amplicon sequencing.

**Results:** High-severity soils reduced seedling growth, EMF colonization, and fungal richness. Low-severity burns supported more rich, diverse fungal communities and greater seedling performance. Stress priming treatments had minimal effects. Fungal community composition varied significantly by burn severity.

**Conclusion:** Fire severity drives changes in fungal community structure that influence seedling outcomes. Low-severity soils retain beneficial microbial legacies, supporting stronger mycorrhizal associations that result in seedling performance comparable to unburned soils. Targeted microbial restoration may enhance post-fire forest regeneration in high-severity burn areas

## Introduction

While recent large-scale wildfires have dominated headlines and understandably drawn widespread public concern, fire has long-served as a natural ecological force – and even an essential ecosystem service – in the development and maintenance of western forests (McKelvey et al. 1996; van Wagtendonk et al. 2018). In addition to shaping vegetation dynamics, fire plays a critical role in soil biogeochemical cycling by altering nutrient availability, soil structure, organic matter inputs, and microbial activity (Certini 2005; McLauchlan et al. 2020; Agbeshie et al. 2022). Historically, however, fire regimes prior to the mid-20th century were characterized by frequent, smaller-scale, mixed-severity events, often with median return intervals under two decades (Stephens et al. 2004; Odion et al. 2014). Such fire regimes promoted the development of fire-adapted plant and microbial communities, contributing to forest structures that were more open, heterogenous, and resilient to disturbance. Frequent, low- to moderate-intensity burns reduce surface fuel loads, limit the encroachment of shade-tolerant species, and maintained conditions favorable to fire-adapted understory vegetation and ectomycorrhizal fungi (EMF) (Fulé et al. 1997; Busse et al. 2000; North et al. 2009; Glassman et al. 2016). Over the past century, fire regimes in the western United States have shifted from these historical patterns to include increasingly frequent, severe, high-intensity megafires that often result in stand-replacing events across extensive landscapes (Parks et al. 2023). Several interacting climatic and anthropogenic factors have contributed to this change in fire regimes. Notable drivers of this transformation include over a century of fire suppression and exclusion, forest densification with associated fuel accumulation, human encroachment on the urban-wildland interface, and the intensification of drought under a warming climate (Neary et al. 1999; MacDonald et al. 2023; Turco et al. 2023; Guo et al. 2024; Kreider et al. 2024). The removal of traditional Indigenous fire stewardship practices, along with legacy land-use changes, have further compounded these effects (Kimmerer and Lake 2001; Safford and Stevens, 2017; Kolden 2019).

As fire severity and frequency shift beyond historical norms, the ecological consequences extend belowground, destabilizing plant-microbe interactions and impairing soil biogeochemical processes critical to post-fire recovery. Fire can have drastic effects on soil physical and chemical properties through the combustion and volatilization of surface organic matter, pH shifts due to ash deposition, and changes in concentrations of key macronutrients like nitrogen and phosphorus. These changes often reduce long-term organic nutrient pools while causing short-term increases in mineral forms (Neary et al. 1999; Certini 2005; Peay et al. 2010; Pellegrini et al. 2020). These changes may either constrain or enhance microbial activity, depending on burn severity, soil type, and post-fire environmental conditions (Fox et al. 2022).

Mutualistic relationships, such as those between forest trees and EMF, may be especially sensitive to changes in fire frequency, severity and seasonality. Successful reestablishment of root symbioses, particularly with EMF, is critical for seedling establishment and performance in post-fire ecosystems (Nara, 2006; Teste et al. 2009; Dove and Hart 2017). This mutualistic relationship influences nutrient uptake, biotic and abiotic stress resistance, and overall biomass accumulation (Hart et al. 2005; Zhang et al. 2024). The modified nutrient profiles and increased pH of burned soils have been shown to disrupt fungal community dynamics and reduce ectomycorrhizal colonization, particularly following high-severity burns (Glassman et al. 2016; Reazin et al. 2016; Pulido-Chavez et al. 2021). While direct soil heating can contribute to microbe mortality near the soil surface, the most significant effects on fungal communities are typically due to indirect changes in soil conditions and plant host availability (Neary et al. 1999; Buscardo et al. 2012; Glassman et al. 2017; Packard et al. 2023). These disturbances reset fungal successional trajectories, often favoring opportunistic or pyrophilous taxa in the early stages, while suppressing more specialized symbionts like EMF until vegetation and soil conditions stabilize (Visser 1995; Glassman et al. 2016; Day et al. 2019; Bruns et al. 2020; Barreiro and Díaz-Raviña 2021; Pulido-Chavez et al. 2021; Packard et al. 2023; Pérez-Izquierdo et al. 2023).

These successional dynamics both above- and below-ground can also be strongly influenced by variation in fire intensity (the energy released during combustion) and severity (the degree of ecological change caused) (Reazin et al. 2016; Smith et al. 2021). While high-severity fires can detrimentally degrade plant and soil biological communities, low-intensity prescribed burns often act as a more controlled disturbance that facilitates activation of dormant plant propagules, minimally affecting soil microbial assemblages, and, in some cases, even enhance microbial richness (Pausas and Lamont, 2022; Fischer et al. 2023; Rafie et al. 2024). These differences in fire behavior are especially relevant when assessing the resilience and recolonization capacity of mutualistic symbionts such as EMF, whose ability to reestablish root associations may depend on the severity of the fire from which they are recovering (Hart et al. 2005; Glassman et al. 2016; Bowman et al. 2021; Nelson et al. 2022). Disruptions to mycorrhizal fungal inoculum caused by severe fires can limit colonization potential, thereby reducing seedling growth even in otherwise favorable soil conditions. Conversely, low-severity burns often preserve fungal networks and soil conditions, helping facilitate faster root colonization and enhanced plant performance (Peay et al. 2009; Peay et al. 2010; Glassman et al. 2016; Dove and Hart 2017).

Drought stress is another major limiting factor in seedling establishment, particularly in post-fire environments where soil structure, organic matter, and microbial communities are often severely disrupted. In recent years, drought-priming techniques–such as the application of mild water stress or phytohormone signalling compounds like abscisic acid (ABA)–have shown promise in enhancing plant tolerance to subsequent water limitation. These strategies aim to preemptively activate physiological and molecular stress-response mechanisms, potentially improving seedling resilience during dry periods (Jacques et al. 2021). However, the effectiveness of such techniques can be influenced by soil microbial communities, which themselves are shaped by abiotic stress, including subsequent post-fire successional processes (Canarini et al. 2021).

Understanding how drought priming interacts with microbial legacies across burn gradients is critical for developing scalable reforestation practices amid increasingly variable climate conditions.

Despite the growing recognition of the role EMF play in forest recovery, relatively little is known about how fire severity and post-disturbance soil conditions jointly influence the reestablishment of symbioses and seedling performance. While previous studies have documented fire-driven shifts in fungal community composition and soil chemistry, fewer have linked these biotic and abiotic changes to root colonization outcomes or tested how microbial legacies interact with plant physiological treatments. To address these gaps, we conducted a greenhouse bioassay using soils collected from forest sites spanning a gradient of fire severity. We investigated how fire history, associated changes in soil chemical properties, and a drought priming treatment using exogenous ABA influenced fungal community structure, EMF colonization, and Douglas-fir (*Pseudotsuga menziesii*) seedling growth. We hypothesized that (1) fungal communities in high-severity burned soils would be compositionally distinct and dominated by ruderal taxa relative to those in low-severity and unburned soils; (2) EMF colonization and seedling biomass would be reduced in high-severity soils due to microbial and chemical disruption; and (3) ABA priming would improve drought tolerance and growth across treatments. This study aims to advance the understanding of how shifting fire-regimes, soil abiotic legacies, and nursery-based physiological interventions shape belowground mutualisms and early-stage seedling performance in post-fire forest ecosystems.

## Materials and Methods

### Study Site

We collected soil inocula on June 5, 2023, at Blodgett Forest Research Station, which sits amid a mixed-conifer forest in El Dorado County, CA approximately 115 km east of Sacramento, CA. Dominant conifer species consist of *Pseudotsuga menziesii*, *Pinus ponderosa*, *Pinus lambertiana, Abies concolor*, and *Calocedrus decurrens*. The most common hardwood species is *Quercus kelloggii*, with *Arbutus menziesii, Notholithocarpus densiflorus, Cornus nuttallii, Alnus spp.* and *Acer spp.* being found near streams. This 1763 ha site in the Sierra Nevada Mountains lies at elevations between 1188 m and 1463 m above sea level. The climate for El Dorado County is chiefly Mediterranean. Annual temperatures for the Blodgett facility range from as low as 0°C in the winter to 32°C in the summer. Mean annual precipitation for the sampling area is 1651 mm and may range from 570 to 2740 mm. Snowfall averages 2540 mm annually. The parent materials of the major soil series–Holland, Musick, and Piliken-variant–are granodiorite, with Cohasset soils forming from volcanic andesite. The natural fire regime is dominated by low-severity fires with high-severity within-stand fires occurring occasionally. Without suppression methods being enacted, the return interval for fire disturbance is 13 years (Berkley Forests 2024).

We identified four different sites which had experienced fires of varying severity at different times. Two sites were subject to low severity prescribed burns as part of the Fire and Fire Surrogate Study that began in 2001 (Stephens and Moghaddas 2005). The first of these sites experienced low severity prescribed burns in 2002, 2009 and 2017. The remaining low severity sampling site was burned in Spring 2020 and one week before sampling in May 2023. The high severity burn site was amid the Mosquito Fire burn scar and last burned in October 2022 at sample collection. Trees in this area exhibited extensive trunk charring and high mortality rates. The final sampling site was noted in Blodgett records to have not experienced a fire in over 100 years and had a thick duff layer measuring from 15-27 cm.

### Soil Sampling and Preparation

At each of the four sites, two living Douglas fir (*Pseudotsuga menziesii*) and two living *Pinus ponderosa* were identified from under which to collect soil samples. We removed the duff layer from an area within 1 m of the tree before collecting soil between 30 cm and 60 cm from the base of the trunk, to a depth of approximately 15 cm. Using a 70% ethanol-sterilized shovel, soil was carefully dug and placed directly into a sterile 2 L whirl-top bag. Each bag of soil was immediately placed on ice in an ice chest for transport back to the University of California, Davis.

Samples for each tree type at each location were passed through a 4 mm sieve and pooled together, with a subsample of each pool reserved for pH and moisture measurements.Half of each soil inoculum was autoclave sterilized (121°C for 45 minutes) twice, separated by 24 hrs, to provide a control group for each treatment. Prior to planting, the soil treatments were mixed with Quikrete playground sand (Atlanta, GA, USA), sterilized in the same manner as the soil, resulting in a 1:1 soil-sand mixture. In all, there were 8 soil inoculum treatments based on burn severity and time since burn: Low (Old), Low (New), High, and Never, with each containing a sterilized and unsterilized soil treatment.

We measured soil pH for a portion of the reserved subsample of each pooled soil within 2 days in a 0.01 M CaCl_2_ solution (Ryti 1965) with an Orion Star 8211 pH-sensitive electrode (Thermo Fisher Scientific, Waltham, MA, USA). To quantify soil moisture and convert between fresh, air dried, and oven dried masses, we air dried 100g of soil for 72 hrs, took the mass, then oven dried the same sample at 60°C for 24 hours and weighed it within 30 mins of removal from the oven.

Other soil chemical analyses were conducted by Brookside Labs (New Bremen, OH) who performed standard soil chemistry tests (including Bray II phosphorus and total nitrogen), carbon to nitrogen ratio (C:N), and soil texture.

### Bioassay Preparation and Initial Growth Phase

*Pseudotsuga menziesii* seeds collected in Oregon, USA, were procured from Sheffield’s Seed Co. (Locke, NY, USA) and surface sterilized in 4.5% H_2_O_2_ with a drop of Tween 20 for 20 minutes. After being rinsed with sterile MilliQ water, the seeds were transferred to a sterile beaker containing MilliQ water and a magnetic stir bar, and left to stir for 24 hours. To stratify, seeds were placed on sterile water agar, sealed with parafilm, and left at 4°C for 28 days.

We conducted our bioassay in a temperature-controlled greenhouse to best mimic the natural growth conditions of Douglas-fir in the Sierra Nevada. To set up the greenhouse experiment, 164 mL Cone-tainers (cones) (Stuewe and Sons, Tangent, OR, USA) were sterilized in 10% bleach solution (BradyPlus, Waxie Sanitary Supply, San Diego, CA), and thoroughly rinsed with deionized (DI) water. Labeled cones were placed in racks and grouped by their soil type and sterilization status. Planting occurred on 26 July, 2023. Each cone was lined at the bottom with Poly-fil polyester fiber (Fairfield Processing Corporation, St. Louis, MO, USA) and filled with the 1:1 soil-sand mixture up to approximately 2 cm below the top rim, and then saturated with reverse-osmosis water. Using 70% ethanol-sterilized forceps, a shallow hole approximately 5 mm deep was made in the soil. Seeds were placed in the hole and covered with soil. A single seed was planted in 331 cones, and 234 cones were planted with two seeds in case of germination failure. In all, 565 cones contained 799 seeds. After planting, each rack was covered in Saran Wrap to maintain humidity prior to setting up irrigation. For the germination phase of the experiment, the racks were arranged in three rows on a single greenhouse bench. Cones were randomly, but evenly, distributed among the 15 racks to account for potential variation in greenhouse conditions. The cones were placed in 96-slot racks, filling every other slot, totaling no more than 48 cones per rack. During the germination and initial growth phase, each cone received RO water through drip irrigation from stake emitters twice a day for one minute, programmed at a rate of 16.67 mL/minute. Seedlings were checked daily to track germination and to remove any dead or potentially pathogen-infected seedlings. All viable seedlings had visible cotyledons emerging on 20 September, 2023, signalling the end of the germination and initial growth phase.

### Stress Priming Treatments

To ascertain the impact of stress priming on Douglas-fir seedling and EMF response to acute drought, we implemented stress priming treatments via water limitation and ABA application, starting four weeks prior to an acute dry-down, during which no water was applied. Prior to the implementation of the priming treatments, we learned that the irrigation system had not been emitting even volumes of water. We therefore switched to hand watering with an Eppendorf repeater pipette 3-4 times per week. For the priming treatments, the seedlings were broken up into three groups, with each group on three different tables to reduce treatment errors. The water limitation group received 4 mL of water every other day; the ABA treatment group received 10 mL of an ABA solution once weekly, and 10-15 mL of water at the other watering sessions; and the control group received 10-15 mL water (water to saturation) at the same interval. The ABA treatments were further divided among three experimental concentrations: 40 seedlings received a 100 µM concentration, 46 received 300 µM, and another 46 received 500 µM ABA. Due to the limited number of surviving seedlings planted into high severity, sterilized soils, that group was omitted from the 100 µM treatment. Soil moisture was regularly tracked using a handheld moisture monitor. This ensured the reduced water group remained below 15% soil moisture to induce a drought response, and that the two normal water groups stayed above this threshold. After the ABA treatments had concluded, watering was stopped entirely for three weeks to dry down the plants immediately prior to harvest, which occurred on 28 March 2024.

Just before harvest, to assess the seedlings’ responses to the priming treatments, we used a PAM-2000 fluorometer (Heinz Walz GmbH, Effeltrich, Germany) to measure the dark-adapted chlorophyll fluorescence (Fv/Fm) of 146 plants, including all burn severity and stress priming treatment groups. On the day measurements were taken, the sun was fully set by 9:00 pm. Readings were taken over two days after 9:30 pm to ensure the plants were dark-adapted.

### Harvest and Post-Harvest Data Collection

After 9 months of growth, the seedlings were harvested.The leaf color for each shoot was graded on a scale from 1 to 5 (1 being completely yellowed and 5 being vibrant green), recorded, and the shoot placed in a labeled plastic bag. Fresh shoot biomass was measured using an XSR105 microbalance (Mettler Toledo, Columbus, OH, USA) with the precision of ±0.00001 g. Each shoot was placed in a coin envelope and dried in a drying oven at 60° C to constant mass for 48-72 hours. After drying, dry biomass was measured in the same manner as fresh biomass.

Percent colonization data collection of the root systems was collected in the days following the harvest. Using sterilized scissors, each root system was cut into 1-2 cm fragments, and placed into a petri dish containing DI water. The root pieces were swirled to homogenize and a random selection was placed evenly in a gridded dish. Using a dissecting microscope, the root tips were analyzed for ectomycorrhizal colonization. At least 200 total root tips - colonized and uncolonized - were counted per seedling, tracking the total terminal roots analyzed with a multi-button clicker counter. The criteria for identifying colonized roots was, (1) a visible mat of hyphae is required, (2) root hairs must be reduced on at least part of the root tip, (3) root tip may potentially be swollen, (4) unusual branching may be present, (5) coralloid clusters are counted as a single root tip, (6) a poorly developed mantle should not be counted as colonized. (Pérez-Pazos et al. 2021). Once ≥200 root tips had been counted, the totals on the clicker counter were recorded. For each root system counted, 8 random colonized root tips were removed with sterile forceps and placed into 20 µL of DNA extraction solution and stored at 20°C for future molecular analysis. To determine percent colonization, the number of colonized tips was divided by the total number counted. Upon completion of a root system, the root fragments were patted dry on a Kimwipe. The fresh biomass of the root sample was measured, placed in a drying envelope, and dried in the same manner as the shoots before recording dry biomass on the microbalance.

### DNA Extraction, Sanger Sequencing and Fungal Identification

We prepared a CTAB-based root-tip extraction solution adapted from methods developed in the Vilgalys Lab (unpublished protocol). Briefly, in a clean 100 mL vessel we combined 10 mL of 1 M Tris-HCl (pH 8.0), 1.86 g KCl, and 0.37 g EDTA, added 80 mL deionized water, adjusted the pH to 9.5–10 with 1 M NaOH, and brought the volume to 100 mL; colonized root tips were placed in 20 µL of this solution. The tips were placed in a thermal cycler and incubated at 65°C for 10 minutes, and then 95°C for 10 minutes. After removing the samples from the thermal cycler, 20 µL of filter-sterilized 3% BSA dilution solution (MilliporeSigma, Burlington, MA, USA) was added to each tube, and samples were stored at -20°C.

In total, a subset of 50 seedlings had the DNA extracted, amplified and sequenced on 8 colonized root tips. All burn treatments were represented, with only the low water and full water treatments included. We omitted the ABA treatment group from fungal community analysis.

Although most seedlings in sterilized soil were not colonized, 5 of these had significant colonization (40% or more), so were included in molecular fungal community characterization. The fungal internal transcribed spacer region (ITS) was used for DNA amplification with ITS1F forward and ITS4 reverse primers (IDT, Coralville, IA, USA) (White et al. 1990; Gardes and Bruns 1993), respectively. Polymerase chain reaction (PCR) amplification was performed using 10 µL GoTaq Green Master Mix (Promega, Madison, WI, USA), 1 µL of each primer, 7 µL ultrapure water, and 1 µL of extracted DNA per sample. PCRs performed used the following settings: To start, 1 minute at 95°C, then 30 cycles of 94°C for 1 minute, 51°C for 1 minute, and 72°C for 1 minute. After the final cycle was closed, samples were annealed at 72°C for an additional 8 minutes. Samples were transferred to -20°C for storage prior to gel electrophoresis. Gels were prepared using 3% ultrapure agarose gel (Invitrogen, Thermo Fisher Scientific, Waltham, MA, USA) in 1x Tris/Borate/EDTA (TBE) buffer (Thermo Fisher Scientific, Waltham, MA, USA), using GelRed stain (MilliporeSigma, Burlington, MA, USA). 3-5 µL of PCR products were added to each well. For samples with multiple bands, PCRs were redone using the ITS4B reverse primer (Gardes and Bruns 1993). Verified samples were transferred to 96-well plates and sent to MCLAB (South San Francisco, CA) for Sanger sequencing. Chromatograms were analyzed using FinchTV (Geospiza Inc., Seattle, WA). Resulting nucleotide sequences were identified using the BLAST tool provided by the National Center for Biotechnology Information (NCBI). Thresholds for genus and species identification were 90% and 97% match, respectively, with e-values ≤1e-20 and ≥95% query cover. However, to account for variability in sequence quality and limited discriminatory power of ITS within certain genera (e.g., *Rhizopogon*), a relaxed threshold of 95% was applied in some cases. When multiple species matches had similar scores or fell near the 95-97% range, taxa were assigned to the genus level. In cases where one species clearly outmatched others, species-level identification was retained. To avoid over-interpretation, these taxa were visualized accordingly in community composition visualizations (Figures 5, 6 and 7).

## Data Analysis and Statistics

All statistical analyses and visualizations were performed in R (v4.4.1; R Core Team, 2024) using RStudio (v2024.9.1.394). Data manipulation and preprocessing were conducted with the dplyr, tidyr, readr, and stringr packages. Fungal community data were integrated and structured using the phyloseq package. Diversity metrics, Bray-Curtis dissimilarity matrices, non-metric multidimensional scaling (NMDS), and permutational multivariate analysis of variance (PERMANOVA) were calculated using the vegan package. Community dispersion (beta-dispersion) tests were also performed with vegan.

Linear mixed-effects models were fit using lme4 and lmerTest packages. Rack position was initially considered a random effect but did not significantly influence model outcomes and was omitted from final models. Final models included burn severity, stress priming treatment, and soil sterilization as fixed effects, along with their interactions. Response variables were assessed for normality and heteroscedasticity using bptest, shapiro.test, residual plots, and additional tools from the PLS205 custom instructional package (Runcie 2025). Transformations (e.g., arcsin square root) were applied when necessary to meet model assumptions.

Post hoc pairwise comparisons and estimated marginal means were calculated using the emmeans package, with multiple comparisons adjusted using Tukey’s HSD, Dunnett’s test, or Bonferroni correction, where applicable. Compact letter displays were generated using multicompView. Statistical significance was evaluated at α = 0.05 for all tests.

All visualizations were generated with ggplot2. Multi-panel figures were combined and annotated using the patchwork and ggtext packages. Colorblind-friendly HCL palettes were implemented via the colorspace package.

## Results

### The Impacts of Burn Severity on Seedling Success

Burn severity had a significant effect on total dry biomass of Douglas-fir seedlings (ANOVA: F(3, 135) = 6.12, *p* < 0.001). High-severity burned soils significantly reduced biomass compared to low-severity and never-burned soils. Estimated marginal mean biomass was lowest in high-severity soils (mean = 0.248 g, 95% CI: 0.216–0.281) and highest in both low (old) soils (mean = 0.320 g, 95% CI: 0.293–0.346) and never-burned soils (mean = 0.320 g, 95% CI: 0.290–0.351), based on the full linear mixed-effects model including stress priming and sterilization treatments. Biomass in the low (new) treatment fell between these extremes (mean = 0.309 g, 95% CI: 0.280–0.329). Compared to high-severity soils, biomass was significantly higher in never-burned (*p* = 0.005), low (old) (*p* = 0.002), and low (new) soils (*p* = 0.02) (Figure 1).

**Figure 1.**
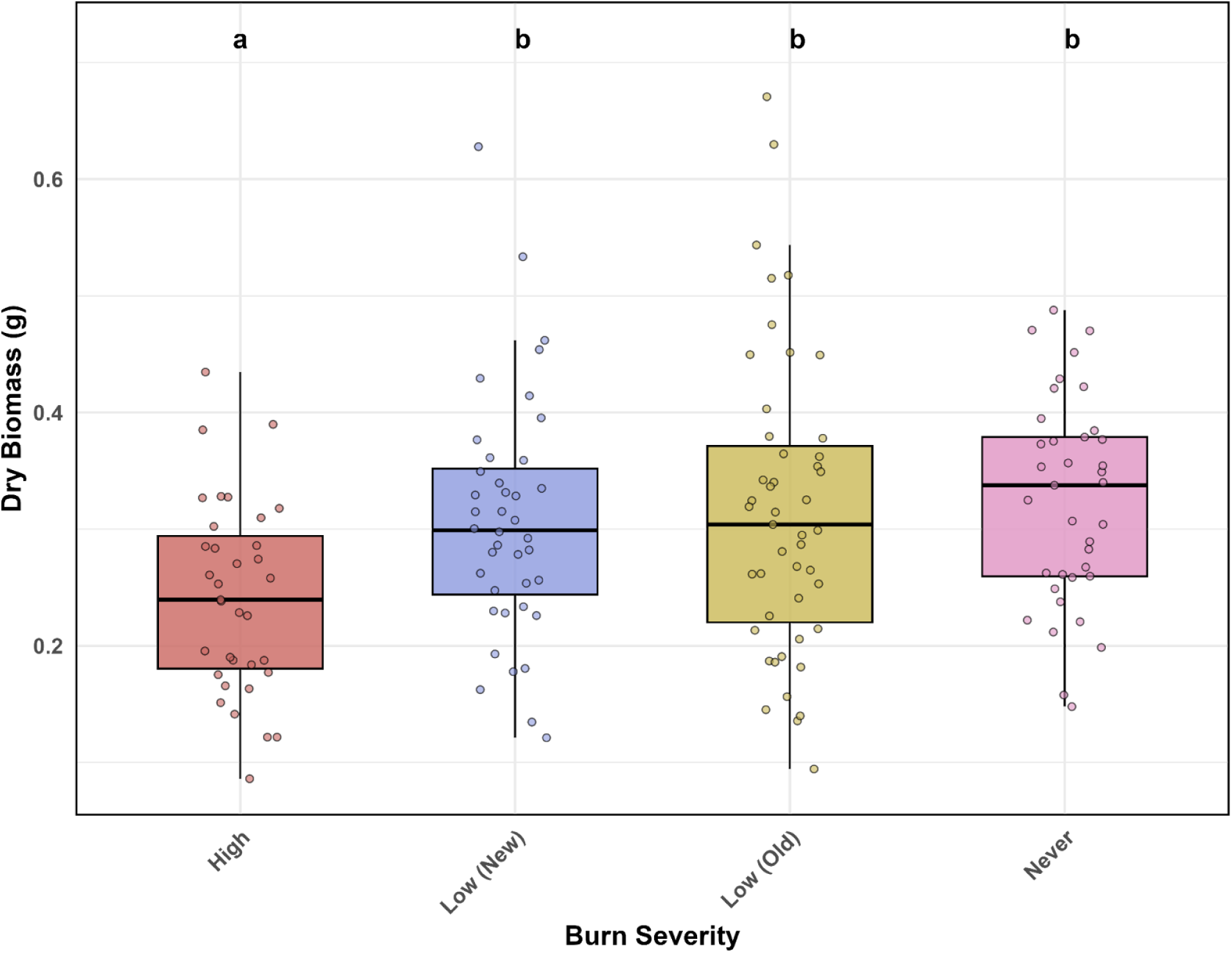
Dry biomass of seedlings grown in soils from four burn severity treatments (n = 160 seedlings total). Each point represents one seedling. Letters above boxplots indicate groupings based on compact letter display (CLD) results from pairwise comparisons of estimated marginal means, adjusted using the Bonferroni method (α = 0.05). Seedlings grown in high-severity soils exhibited the lowest biomass accumulation. No significant differences were observed among Low (New), Low (Old), and Never-burned soils, with Low (Old) and Never treatments showing the highest average biomass.

Although estimated biomass values varied slightly across stress priming treatments, no consistent patterns were observed, and water limitation had no significant effect on biomass, either as a main effect (*p* = 0.84) or in interaction with burn severity or sterilization (all *p* > 0.25). Pairwise comparisons among stress priming treatments also revealed no significant differences (all *p* = 1.00).

Soil sterilization had a significant effect on seedling biomass (*p* < 0.001), and interacted significantly with burn severity (*p* = 0.002). In both low-severity soils, biomass was highest when soils were unsterilized, particularly in low-severity (old) soils (mean = 0.406 g), followed by low-severity (new) soils (mean = 0.348 g). Sterilization substantially reduced biomass in both cases, with means of 0.234 g and 0.271 g, respectively. A similar reduction was observed in never-burned soils, where unsterilized soils supported greater biomass accumulation (mean = 0.344 g) compared to sterilized soils (mean = 0.297 g). In contrast, seedling biomass in high-severity soils remained low regardless of sterilization status (means = 0.236 g and 0.261 g) Model diagnostics confirmed that assumptions of normality and homoscedasticity were reasonably met (Supplemental Figure).

### Percent Colonization

After arcsine square root transformation to meet model assumptions, burn severity significantly affected percent colonization of Douglas-fir seedling roots (ANOVA: *F*(3, 37) = 3.02, *p* = 0.042). Estimated marginal means indicated that colonization was lowest in seedlings grown in high-severity burned soils (34.3%), and highest in those from low-severity (new) (52.3%) and low-severity (old) (50.3%) soils. Mean percent colonization in never-burned soils (41.6%) fell in between the high-severity and low-severity treatments (Figure 2).

**Figure 2.**
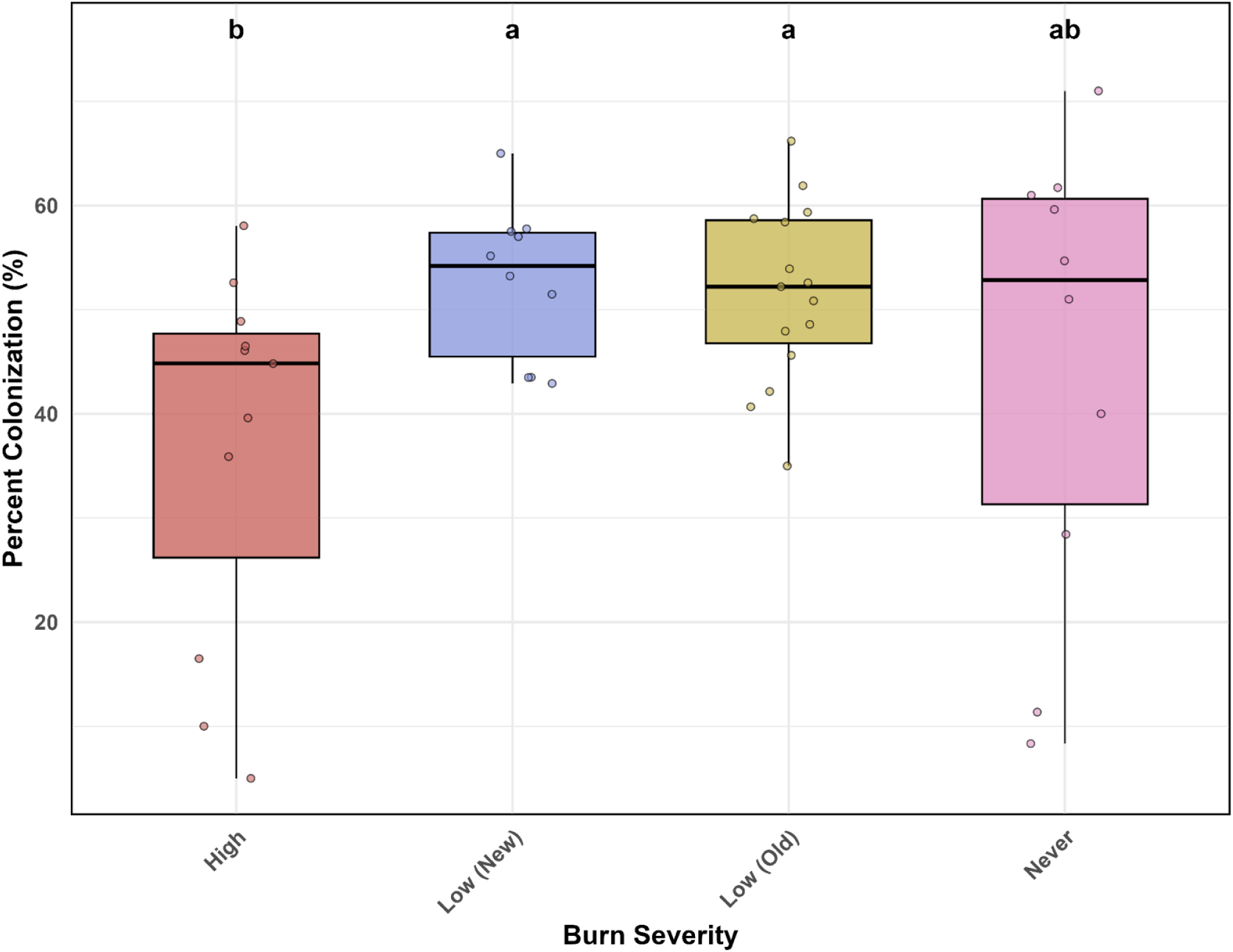
Percent of seedling root tips colonized by ectomycorrhizal fungi across burn severity treatments. Each point represents one seedling. Letters above boxplots indicate significant differences between burn treatments based on a compact letter display from pairwise comparisons of estimated marginal means (Dunnett-adjusted, α = 0.05). Colonization was highest in seedlings grown in low-severity (old) soils, though not significantly higher than those in low-severity (new) soils. Both low-severity soils supported significantly higher colonization than high-severity soils, but were not significantly different from never-burned soils. No significant difference was observed between high-severity and never-burned treatments.

Post-hoc pairwise comparisons using Dunnett’s test revealed that colonization in low-severity (new) soils was significantly higher than in high-severity soils (*p* = 0.042), and colonization in low-severity (old) soils was marginally higher (*p* = 0.053) than low-severity (new) and remained significant. No significant difference was observed between high-severity and never-burned soils (*p* = 0.589), nor between either low severity and never-burned soils.

Stress priming treatments had no significant effect on colonization (*p* = 0.375), nor was a significant interaction observed between priming treatment and burn severity (*p* = 0.195).

### Maximum Quantum Yield of PSII

Photochemical efficiency (measured as Fv/Fm) was analyzed after the acute dry-down to determine the effects of burn severity, soil sterilization, and stress priming treatments - including ABA - on seedling physiological drought responses (Figure 3). Burn severity and soil sterilization significantly influenced Fv/Fm (ANOVA: F(3, 126) = 4.10, *p* = 0.008; F(1, 126) = 175.75, *p* < 0.001, respectively), while stress priming treatments had no significant effect (*p* = 0.75). No significant interactions were observed among treatment factors.

**Figure 3.**
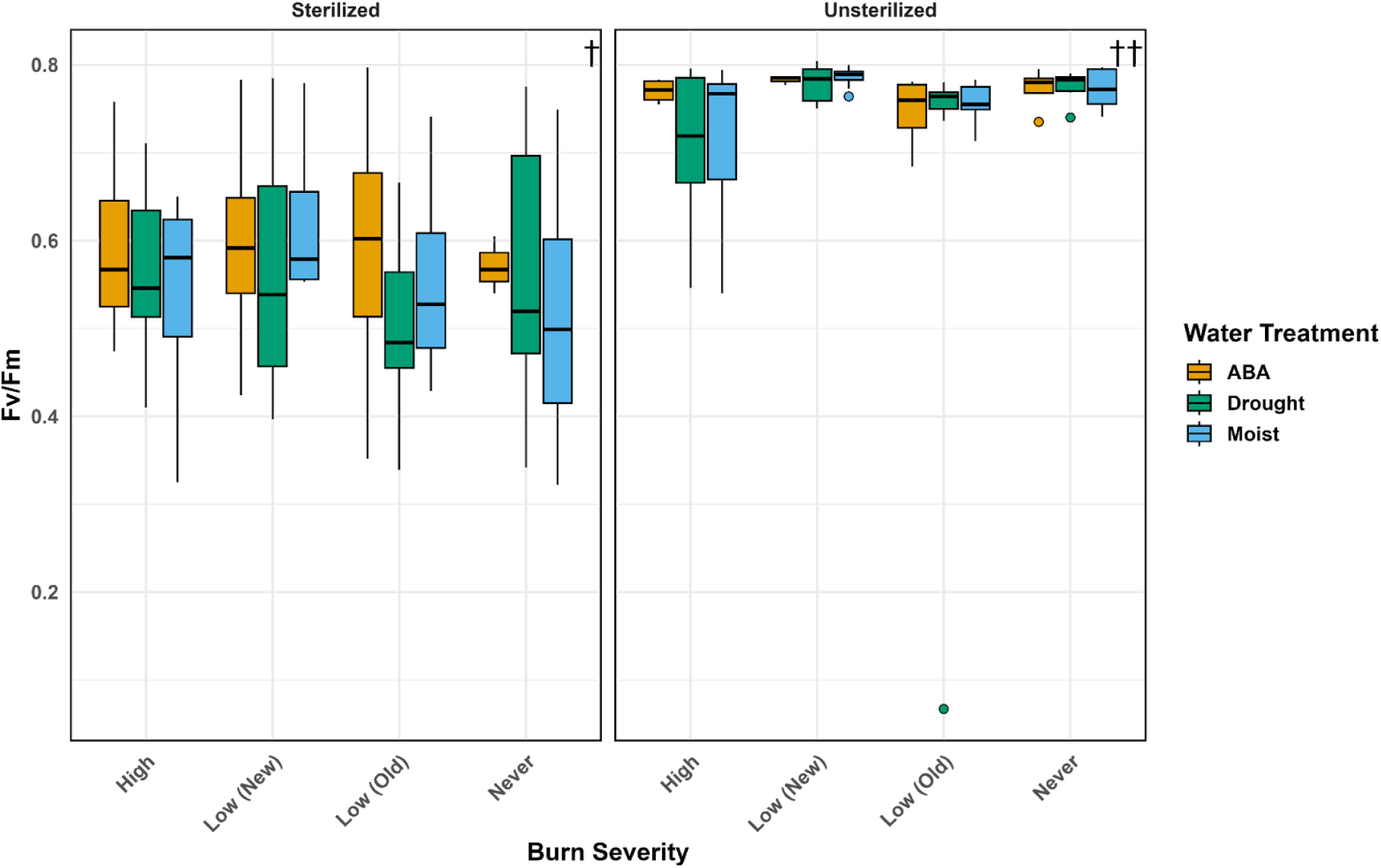
Boxplot of chlorophyll fluorescence (Fv/Fm) in seedlings grown under different burn severities, water treatments, and soil sterilization statuses. All seedlings underwent a three-week dry-down prior to chlorophyll fluorometer measurement. No significant differences in photochemical efficiency were detected among water or burn treatments within each sterilization group. However, significant differences were observed between sterilized and unsterilized soils, emphasizing the role of microbial communities in maintaining photochemical efficiency and supporting seedling performance. († denotes significance between sterilization treatments).

Means estimates indicated that Fv/Fm was lowest in seedlings grown in high-severity, sterilized soils (mean = 0.561, 95% CI: 0.515-0.606) and highest in unsterilized soils from low-severity (new) (mean = 0.783, 95% CI: 0.742–0.821) and never-burned treatments (mean = 0.774, 95% CI: 0.735–0.811). Sterilization had a consistently negative effect on Fv/Fm across all burn severities, with unsterilized soils producing higher Fv/Fm values than their sterilized counterparts. The difference in Fv/Fm between high-severity sterilized and low-severity unsterilized soils was approximately 0.22, representing a ∼39% increase in photochemical efficiency.

Despite the significant overall effect of burn severity, Tukey-adjusted pairwise comparisons revealed no statistically significant differences between individual burn levels (all *p* > 0.19). However, all comparisons between sterilized and unsterilized soils were highly significant (*p* < 0.001).

To evaluate whether photochemical efficiency differed among ABA concentrations, we conducted a subset analysis including only ABA-treated seedlings. ABA concentration (100, 300, or 500 µM) did not significantly affect Fv/Fm (*p* = 0.17). Back-transformed means ranged from 0.63 (100 µM) to 0.71 (500 µM), and pairwise comparisons revealed no significant differences between concentrations (all *p* > 0.36).

### Fungal Community Composition, Structure and Species Richness

#### Species richness

Species richness per seedling varied across burn severity treatments (Figure 4). For both all fungal taxa and the EMF-only subset, seedlings grown in high-severity burned soils generally hosted fewer unique species than those from low-severity and never burned soils. Richness was typically higher in low-severity (old) soils, followed by low-severity (new) and never burned soils. These trends were more pronounced in the EMF dataset, where several seedlings grown in high-severity burned soils lacked any detectable EMF taxa. Species counts were more variable in the all-taxa dataset, particularly for the high- and low-severity treatments, whereas EMF richness exhibited a more uniform distribution across treatments. However, richness values in never-burned soils displayed similar ranges in both all-taxa and EMF-only datasets. Notably, the low-severity (old) treatment contained four possible outliers with six distinct EMF taxa each, representing the highest observed richness in either dataset.

**Figure 4.**
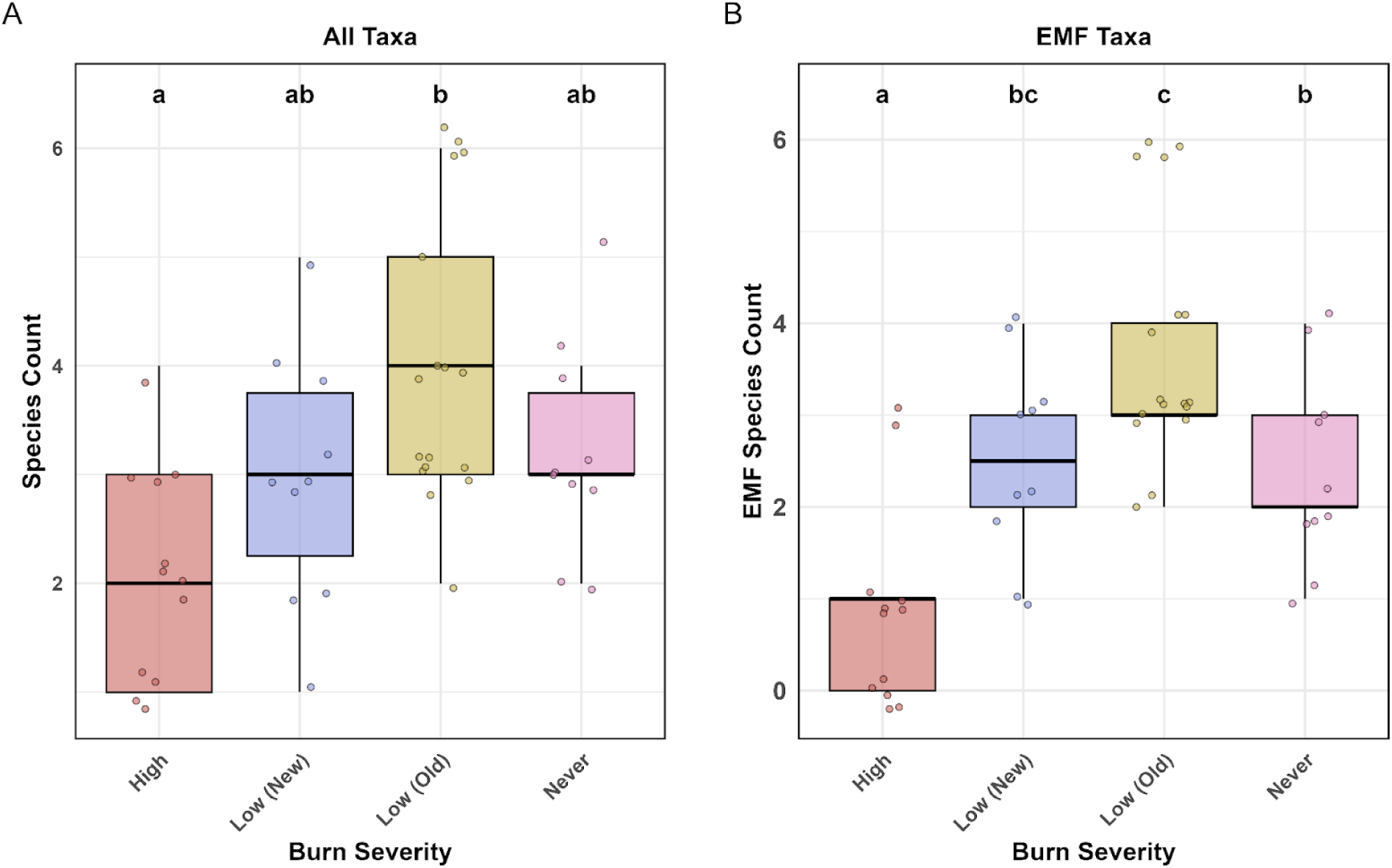
Species richness of fungal communities detected on seedling root tips across burn severity treatments. (A) Boxplot showing richness of all fungal taxa identified via Sanger sequencing from eight root tip samples per seedling. Letters denote significant differences among treatments based on Tukey’s HSD test. Low (Old) soils supported the highest overall richness and were the only treatment significantly different from High-severity soils.(B) Boxplot showing richness of ectomycorrhizal fungi (EMF) after filtering to include only sequences identified as EMF. Both Low-severity treatments and the Never-burned soils supported significantly higher EMF richness than High-severity soils. Low (Old) soils again supported the highest richness and differed significantly from the Never-burned treatment

Our one-way ANOVA verified the significance of burn severity on overall species richness per seedling (ANOVA: *F*(3, 45) = 6.71, *p* < 0.001). Seedlings from high-severity soil had the lowest richness (mean = 2.08, 95% CI: 1.42–2.75), while those from low-severity (old) had the highest (mean = 4.00, 95% CI: 3.44–4.56). Never-burned soils also supported higher richness (mean = 3.20, 95% CI: 2.47–3.93), followed by low-severity (new) (mean = 3.00, 95% CI: 2.27–3.73).

Post-hoc comparisons (Tukey’s HSD) for all taxa confirmed that species richness in high-severity soils was significantly lower than low-severity (old) soils (*p* < 0.001). No other pairwise comparisons reached statistical significance, though richness was lower in high severity soils across all contrasts. Neither the stress priming treatment (*p* = 0.95) nor its interaction with burn severity (*p* = 0.41) had any significant effect and were excluded from further analyses.

Focusing exclusively on EMF taxa present on seedling roots, we applied the same ANOVA and post-hoc comparison methods used in the total fungal richness analysis. This revealed more pronounced differences among burn severity treatments (ANOVA: F(3, 45) = 13.26, *p* < 0.001). Mean EMF species richness was lowest among high-severity soils (mean = 0.92, 95% CI: 0.22–1.61) and highest among low-severity (old) soils (mean = 3.77, 95% CI: 3.18–4.35).

Richness in low-severity (new) soils and never burned soils was intermediate (low-severity (new): mean = 2.50, 95% CI: 1.74–3.26; never burned: mean = 2.40, 95% CI: 1.64–3.16).

Tukey’s HSD post-hoc comparisons indicated that richness was significantly lower in high severity soils compared to low-severity (old) (*p* < 0.001), low-severity (new) (*p* = 0.018), and never burned soils (*p* = 0.029). One marginal contrast between low-severity (old) and low-severity (new) did approach significance (*p* = 0.053), while richness in never burned soils was significantly lower than low-severity (old) soils (*p* = 0.032). The remaining contrast between never burned soils and low-severity (new) soil was not significant (*p* = 1.0).

#### Spatial turnover in species composition

Nonmetric multidimensional scaling (NMDS) ordination for all fungal taxa revealed clear differences in fungal community composition by burn severity (Figure 5). Samples from high severity burned soils were generally separated from those of low-severity and never burned soils along the NMDS1 axis, reflecting compositional shifts. High-severity communities also exhibited greater dispersion, suggesting greater within-treatment variability. In contrast, samples from low-severity and never burned soils clustered more tightly, with considerable overlap between the three groups. These clusters were moderately dispersed along the NMDS2 axis but showed little variation along the NMDS1 axis, indicating broadly similar community structure with some compositional variation.

**Figure 5.**
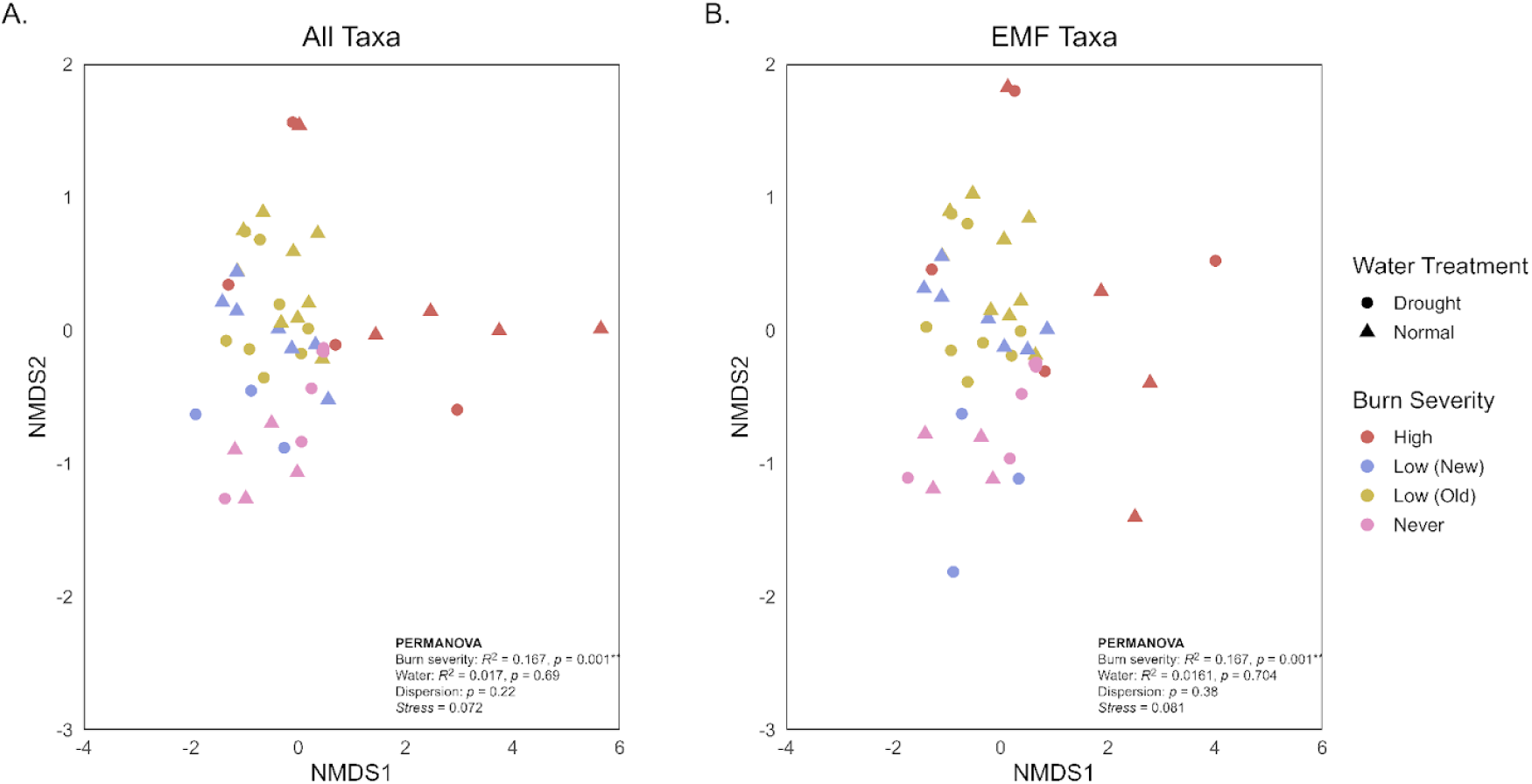
Non-metric multidimensional scaling (NMDS) of fungal community composition in Douglas-fir seedling root tips across burn severity treatments. Each point represents one seedling, colored by burn severity and shaped by water treatment. PERMANOVA results (Bray–Curtis distance) are overlaid on each panel. Water treatment had no significant effect in either ordination. **A**. All Taxa: Fungal communities in high-severity soils formed a distinct cluster with minimal overlap with other treatments. Never-burned and Low (Old) samples also showed relatively tight clustering, while Low (New) samples were more variable and overlapped broadly with other low-severity and unburned groups. **B**. EMF Taxa: Ectomycorrhizal communities showed tighter clustering overall, but with clearer separation among some burn treatments. High-severity samples remained distinct, while Low (New) overlapped with both Low (Old) and Never-burned groups. In contrast, Low (Old) and Never-burned samples showed relatively little overlap with each other, suggesting more differentiated EMF assemblages.

These patterns for overall fungal community composition were validated by PERMANOVA results (Figure 5A). Assessment using Bray-Curtis dissimilarity showed fungal communities differed significantly across burn severity treatment levels (*R^2^* = 0.17, *F* = 2.87, *p* = 0.001). Burn severity alone explained approximately 17% of the variation in fungal community structure. In contrast, stress priming treatment had no significant effect on community composition (*R²* = 0.017, *F* = 0.773, *p* = 0.691). The full model, which included the interaction between burn severity and stress priming treatment, explained 27% of the total variation and remained highly significant (*R²* = 0.29, *F* = 2.28, *p* = 0.001). A test for homogeneity of group dispersions showed no significant differences in within-group variability among burn severity treatments (*F*(3, 43) = 1.53, *p* = 0.220), indicating that the PERMANOVA results reflect true differences in community composition rather than group dispersion.

To determine whether ectomycorrhizal fungi responded similarly to the broader fungal community, we performed a separate NMDS and PERMANOVA restricted to EMF taxa (Figure 5B). The ordination revealed similar patterns: EMF assemblages from high-severity soils were more widely disbursed and distinct from those in low-severity and never burned soils, which showed moderate dispersion and considerable overlap.

PERMANOVA results confirmed that EMF community composition varied significantly across burn treatments (*R²* = 0.16, *F* = 2.66, *p* = 0.001). Burn severity alone explained 16% of the variation in EMF composition, while stress priming treatment had no significant effect (*R²* = 0.020, *F* = 0.86, *p* = 0.550). The full model, including the interaction between burn severity and stress priming, remained significant (*R²* = 0.30, *F* = 2.28, *p* = 0.001). These results indicate that fire severity accounted for the majority of the observed differences in EMF community composition, consistent with patterns observed in the broader fungal community. Variation in community dispersion among burn severity treatments was not significant for EMF taxa (*F*(3, 41) = 0.55, *p* = 0.652).

## Discussion

While fire remains a critical aspect of ecological function, fire severity is a key driver of post-disturbance ecosystem dynamics. As such, the recent shift toward more frequent and severe wildfires presents unique challenges in understanding the belowground responses that shape forest health and resilience in post-fire landscapes. Existing literature has shown that fire severity is a major determinant of seedling success and plays a critical role in restructuring soil microbial communities following disturbance (Glassman et al. 2016; Dove and Hart 2017; Pulido-Chavez et al. 2021). Combined with existing abiotic stressors such as drought, the increased prevalence of high-severity wildfires across the western United States presents a major obstacle for ecosystem stability, forest regeneration, and reforestation. In this study, we examined how a gradient of fire severities influenced the establishment of *P. menziesii* seedlings compared to an undisturbed soil, with a particular focus on the formation of ectomycorrhizal mutualisms essential for seedling survival and growth. Previous work has shown that ectomycorrhizal colonization in burned soils can significantly enhance seedling biomass when compared to sterilized soils (Peay et al. 2009; Jenkins et al. 2018). We therefore aimed to quantify how variation in fire severity affected soil fungal dynamics and seedling performance, and to determine whether stress priming treatments with drought and ABA could improve seedling stress tolerance.

We observed that fire severity had strong and consistent effects on both fungal community structure and seedling outcomes. Seedlings grown in high-severity soil exhibited significantly lower biomass (Figure 1), reduced EMF colonization, and the lowest species richness of both total fungi and EMF relative to other treatments. In contrast, low-severity and unburned treatments supported more diverse fungal communities and higher rates of colonization, which corresponded to greater seedling biomass. Despite the addition of stress priming and ABA treatments to evaluate the potential for physiological priming, neither had significant effects on biomass, colonization, or seedling drought tolerance. These findings underscore the role of fire severity in shaping microbial legacies and early seedling establishment.

Our first hypothesis, stating that fungal communities in high-severity burned soils would be compositionally distinct and dominated by ruderal and stress-tolerant taxa, was supported by both alpha and beta diversity metrics. High-severity soils hosted fungal communities that were significantly less species-rich, particularly in EMF taxa, and showed clear compositional divergence from low-severity and never-burned treatments in NMDS ordinations. Seedlings grown in high-severity soils had significantly lower EMF richness and colonization rates, with some lacking EMF altogether. These findings align with previous studies documenting the loss of fungal diversity and disruption of mutualistic symbionts following severe wildfire (Glassman et al. 2016; Pulido-Chavez et al. 2021; Nelson et al. 2022, Packard et al. 2023; Philpott et al. 2025). These patterns are consistent with post-fire fungal succession where high-severity fire resets the community to an early-successional state dominated by fast-colonizing ruderal pyrophilous taxa. Whereas low-severity and unburned soils maintain more mature fungal assemblages, including late-successional EMF, which may reflect legacy effects or greater inoculum availability (Salo and Kouki, 2018; Kouki and Salo, 2020; Gill et al. 2022).

Taxonomic analysis of root-associated fungi from high-severity seedlings revealed a functionally mixed assemblage, consisting of pyrophilous taxa (52%), along with EMF (29%) and putative pathogens (14%). Among these, *Pustularia–*a known pyrophilous Ascomycete–was the most abundant single taxon with 18 sequences from morphologically colonized ectomycorrhizal root tips. While *Rhizopogon* was the most abundant EMF genus overall, a single *Rhizopogon* species, most closely matching *R. hawkerae* within the *R. vinicolor* clade, accounted for 6 of those, equal to *Myxomphalia maura* (a pyrophilous saprotroph) and *Paraphoma fimeti* (a potential pathogen) (Figure 6). This pattern supports the hypothesis that high-severity wildfire selects for ruderal saprotrophs with some persistence by EMF and pathogenic fungi. The resulting community reflects a functionally mixed, early-successional post-fire assemblage, consistent with prior findings in fire-adapted forest soils (Day et al. 2019; Bruns et al. 2020; Hughes et al. 2020; Smith et al. 2021; Filialuna and Cripps 2021; Pérez-Izquierdo et al. 2023). In contrast, the fungal communities on low-severity and never-burned soils were overwhelmingly dominated by EMF, particularly *Rhizopogon spp.* In never-burned soils, 77% of colonized root tips belonged to the *Rhizopogon* genus, with low-severity (old) and low-severity (new) exhibiting similarly high proportions at 70% and 69%, respectively (Figure 7). This strong dominance by a well-known EMF genus likely reflects both the preservation of fungal inoculum and favorable soil conditions in the less severely burned and never burned environments. Such observations support the idea that lower-severity fire preserves functional EMF networks, enabling greater root colonization and more consistent community structure across seedlings.

**Figure 6.**
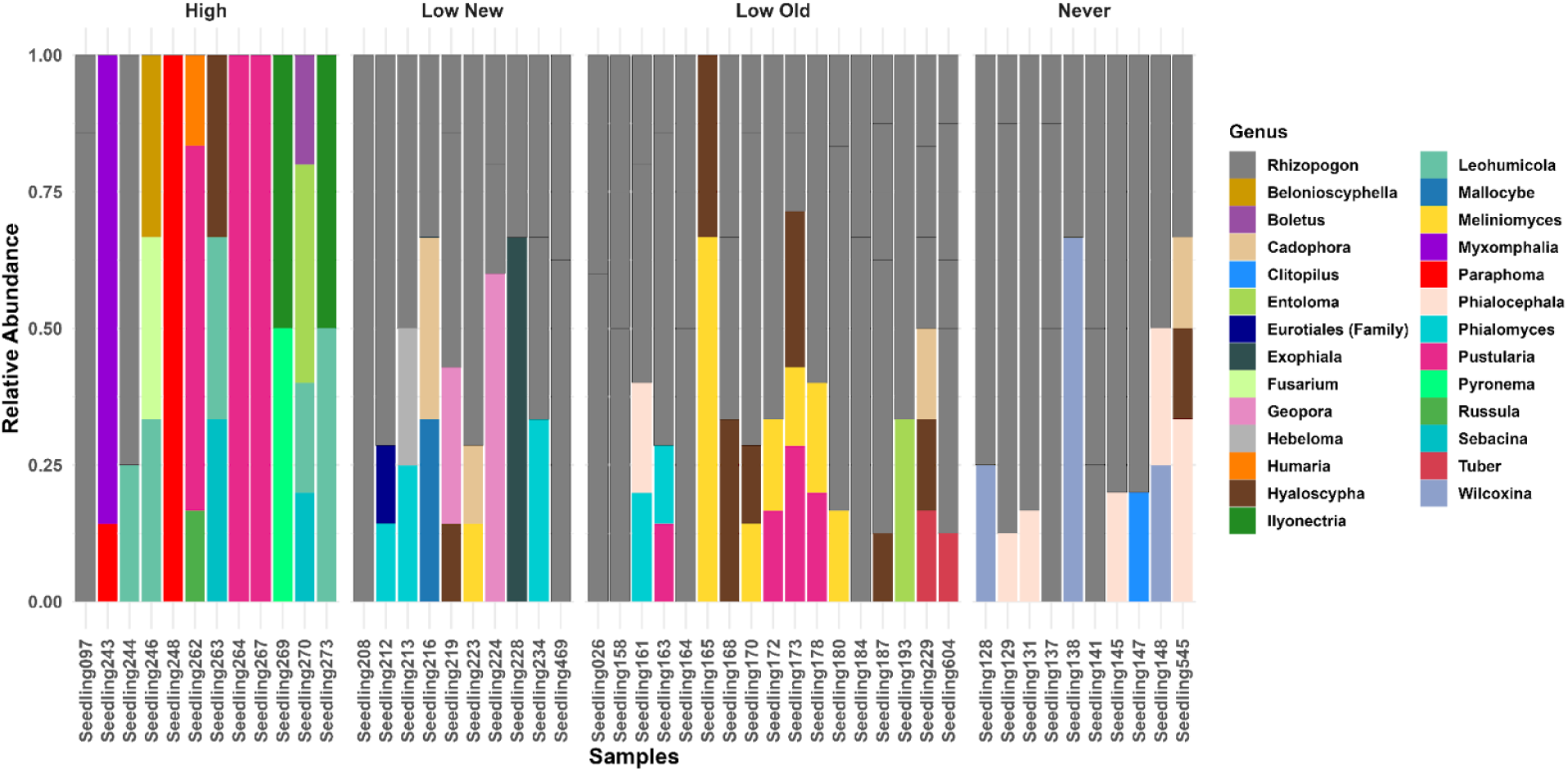
Stacked bar chart showing relative abundance of fungal taxa present on Douglas-fir seedling root tips at the genus level, grouped by burn severity. Grey bars represent Rhizopogon spp., with hash marks indicating species-level differentiation within the genus. Rhizopogon dominated the communities in Low-Severity and Never-Burned treatments. High-Severity seedlings exhibited more variable community composition, including increased relative abundance of pyrophilous saprotrophs and pathogenic fungi.

**Figure 7.**
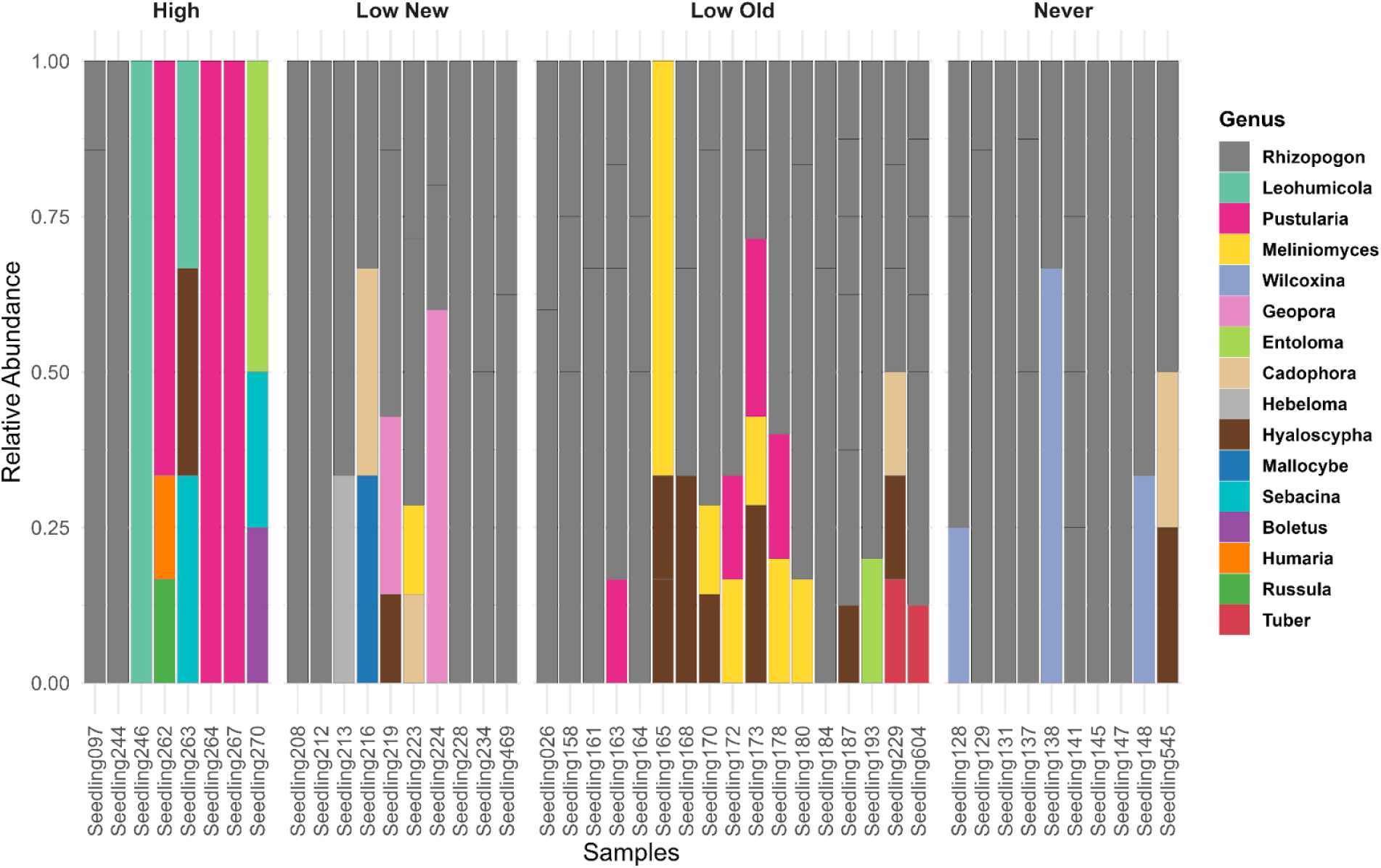
Stacked bar chart showing relative abundance of ectomycorrhizal fungal taxa present on Douglas-fir seedling root tips at the genus level. Rhizopogon again dominated EMF communities in Low-Severity and Never-Burned treatments. High-Severity seedlings supported a more disparate EMF community, with reduced Rhizopogon abundance and greater taxonomic variability. Grey bars with hash marks again represent species-level differentiation within Rhizopogon.

Due to limited discriminatory power in the ITS region for some *Rhizopogon* species and the variable quality of root-derived sequences, several sequences returned high-scoring BLAST matches to both *R. hawkerae* and *R. defectus*, with only marginally lower similarity scores to *R. vinicolor*. Given this ambiguity, we treated these as putative species-level assignments for comparative purposes, while treating ecological patterns at the genus or clade level.

High-severity soils exhibited a trend toward greater within-treatment variability in community composition–although differences in dispersion were not statistically significant (betadisper ANOVA, *p* = 0.38)–suggesting that fungal communities in these environments were not only compositionally distinct but more variable, consistent with a post-disturbance assemblage dominated by opportunistic or ruderal species (Bruns et al. 2020; Packard et al. 2023).

Conversely, low-severity and unburned soils supported more consistent and species-rich fungal communities, including higher EMF richness, suggesting greater legacy retention of symbiotic networks. Although definitive conclusions are limited by sampling resolution, the observed richness trends–with the lowest diversity in the high-severity soils and highest in low-severity (old) soils–are consistent with expectations of the Intermediate Disturbance Hypothesis, which proposes peak diversity under moderate levels of disturbance (Connell 1978). At the community level, however, a small but significant burn x water treatment (drought, full water) interaction indicates that our watering schedule subtly modulated burn-related compositional differences, even though water treatment alone was not a main driver. Collectively, these findings suggest that high-severity fires can reset fungal successional dynamics, favoring disturbance-adapted taxa while constraining the establishment of functionally important mutualists, while low-severity fires have the potential to support increased fungal diversity and community structure.

Both biomass and colonization data supported our second hypothesis: that severely burned soil would reduce ectomycorrhizal colonization and seedling performance. Ectomycorrhizal colonization declined significantly in the high-severity soils, with several seedlings showing minimal or no detectable colonization. However, while analysis showed burn severity to be a significant determinant of percent colonization, the only significant difference shown in post-hoc pairwise analyses was between high-severity and low-severity (new) soils. But although only this single contrast reached statistical significance, both low-severity treatments exhibited markedly higher colonization than the high severity soils, with relative increases of 47-53%.

These patterns, despite limited post-hoc significance, suggest a biologically meaningful reduction in EMF colonization following a high-severity fire. The lack of a significant difference between high-severity and never-burned soils was unexpected, but may reflect differences in fungal recruitment strategies, or more favorable post-fire soil chemistry, than inoculum potential per se. Specifically, colonization in never-burned soils may be lower because EMF taxa in these late-successional communities rely more on stable, established hyphal networks in situ, and may produce fewer or less competitive propagules capable of initiating colonization following soil disturbance. Sampling and subsequent homogenization of soil likely disrupted existing mycelial connections, placing taxa adapted to intact forest systems at a disadvantage relative to disturbance-adapted taxa found in recently burned soils (Glassman et al. 2016; Ishida et al. 2008; Peay et al. 2009).

This pattern may also reflect ecological differences in fungal community composition and successional stage. As previously discussed, low severity soils—particularly the recent burn—were dominated by fire-responsive EMF such as *Rhizopogon*, which have been shown to rapidly colonize seedlings after a disturbance (Kjøller and Bruns, 2003; Peay et al. 2009; Glassman et al. 2015; Van Dorp et al. 2020). By comparison, although *Rhizopogon spp.* were also abundant in never-burned soils, the lower colonization observed could potentially be due to a reliance on established hyphal networks rather than dispersive propagules. While the mild disturbance in low-severity soils may have activated spore banks or increased propagule availability, facilitating more rapid colonization of seedlings (Glassman et al. 2016; Day et al. 2019). The relatively low colonization observed in high-severity soils likely reflects the compounded effects of microbial mortality, loss of viable inoculum, and altered soil physical and chemical properties following severe disturbance (Neary et al. 1999; Dove and Hart 2017; Nelson et al. 2022).

Together, these results suggest that not only burn severity, but also time since fire, fungal response to disturbance, and successional dynamics of EMF communities shape colonization potential. Low-severity burns may create conditions that favor the retention and rapid activity of EMF propagules, supporting higher colonization despite recent disturbance. While both severely burned and long-undisturbed soils may lack the disturbance-adapted fungal traits that support early colonization, resulting in lower establishment of symbioses under greenhouse conditions.

The observed differences in seedling biomass across burn severities were closely linked to the patterns seen in EMF colonization. Seedlings grown in high-severity soils, which supported the lowest levels of colonization and species richness, also exhibited the greatest reductions in total dry biomass. In contrast, biomass in low-severity and never burned soils was approximately 25-30% higher, aligning with more robust EMF communities in those treatments. These patterns support the well-documented role EMF play in nutrient and water acquisition under stressful conditions (Pena and Polle 2014; Chen et al. 2021; Castaño et al. 2023; Sevanto et al. 2023).

EMF not only improve seedling access to key nutrients such as nitrogen and phosphorus, but colonization leads to the formation of extraradical mycelium, effectively increasing root surface area, enhancing water and nutrient uptake outside existing depletion zones. This fungal network extends well beyond the rhizosphere, functioning as an extension of the root system, and allowing for greater soil exploration and access to otherwise unavailable resources. In resource-limited environments, expanding a seedling’s foraging capacity can be crucial for survival and growth (Smith and Read 2008). The loss of essential symbionts in the high-severity soils, as evidenced by both lower colonization rates and reduced species richness, likely limited nutrient acquisition, leading to reduced biomass accumulation. Meanwhile, seedlings in low-severity and never burned soils–where diverse and abundant EMF communities persisted–saw significantly greater biomass accumulation, reinforcing the ecological importance of mutualistic fungi in early seedling development.

In addition to biotic factors, the reduction of seedling biomass in the high-severity burn treatment may also be attributed to multiple interacting biogeochemical processes that impact the soil directly. Wildfires combust surface organic matter, reducing essential carbon sources necessary for microbial activity and nutrient cycling (Certini 2005; Dove et al. 2020; Walker et al. 2019).

Volatilization of nitrogen may also occur during high-intensity burns, creating sustained nitrogen limitation in soils (Neary et al. 1999; Pellegrini et al. 2020). Additionally, heating of surface soil to temperatures between 176°C and 204°C volatilizes hydrophobic substances, which then condense deeper into the soil profile, coating mineral particles and creating a water-repellant layer (DeBano 1981). The creation of this intensely hydrophobic layer reduces water infiltration, further inhibiting plant establishment (Madsen et al. 2016). Finally, the destruction and alteration of soil microbial communities, including EMF, impairs nutrient acquisition and plant development (Dove and Hart 2017; Nelson et al. 2022).

Our soil chemical analyses remained consistent with most of these established patterns, revealing substantial differences between burn treatments. The high-severity soil had notably lower carbon and nitrogen concentrations relative to the low-severity (new) and never burned soils, along with higher pH and increased concentrations of base cations such as calcium and magnesium. The alkaline pH and high base saturation of the high-severity soils was likely driven by ash deposition, leading to possible reduction in the uptake of zinc and iron ions, along with a slight phosphorus limitation due to precipitation of calcium-phosphate minerals. In addition, nitrate concentrations remained low, supporting possible nitrogen limitation contributing to the reduction in seedling biomass in this treatment. Losses of soil organic matter in these soils likely caused a decline in soil structure, water retention, and microbial habitat, further compounding the stresses on seedling establishment. Reduced organic matter measurements in high-severity soils - likely due to combustion and volatilization - supports our direct observation of strong water-repellency, as irrigation water frequently pooled on the surface and was slow to infiltrate during the growth portion of the bioassay. Conversely, low-severity (new) and never burned soils maintained much higher carbon, nitrogen and organic matter levels, water infiltration, as well as pH levels more ideal for plant nutrient uptake, supporting greater biomass accumulation.

Interestingly, the low-severity (old) soil showed reduced carbon, nitrogen, organic matter, and cation exchange capacity compared to low-severity (new) and never burned soils, yet accumulated biomass levels remained remarkably similar. This suggests that soil chemical fertility itself does not fully determine seedling establishment outcomes. And although reduced organic matter in these soils would typically be expected to weaken soil aggregation and water infiltration, the preservation of overall soil structure and retention of biological communities likely helped to offset these effects.

Photochemical efficiency (Fv/Fm) was unaffected by stress priming treatments, providing no support for our third hypothesis that stress priming, whether through ABA application or withholding water, would improve physiological resistance to drought. Instead, Fv/Fm was most strongly affected by soil sterilization, with seedlings in the unsterilized soils showing higher photochemical efficiency across all burn severities. The lowest Fv/Fm values were seen in the high-severity, sterilized soils, while the highest occurred in the unsterilized low-severity and never-burned soils. While burn severity had a significant overall effect, the lack of significant pairwise differences between individual burn levels suggests that the presence or absence of microbial communities–rather than fire history alone–was the primary driver in Fv/Fm variation. These results emphasize the importance of soil biota in regulating plant physiological stress responses and indicate that microbial community integrity may play a more central role than short-term stress priming in supporting seedling function under these experimental conditions. However, it is possible that the duration and intensity of the water limitation and ABA exposure was insufficient to induce a measurable stress priming response.

Our results indicate that high-severity fire can significantly impair EMF colonization, fungal richness, and Douglas-Fir seedling biomass, while selecting for distinct fungal communities favoring saprotrophs–especially the pyrophilous taxa *Pustularia*–and reducing the presence of EMF taxa. Colonization patterns were closely linked to seedling performance, with disturbance-adapted EMF taxa dominating low-severity burned soils and facilitating increased biomass accumulation. In contrast, never-burned soils, while rich in EMF taxa, exhibited lower colonization potential under greenhouse conditions, likely because dominant taxa in these communities depend on established hyphal networks and are less capable of colonizing from spores or hyphal fragments following soil disturbance. Photochemical efficiency was unaffected by stress priming, but soil sterilization consistently reduced Fv/Fm, highlighting the importance of microbial presence in mitigating physiological stress. However, a more extended priming treatment and prolonged subsequent dry-down period may be more effective at eliciting a treatment effect. Importantly, these findings show that low-severity fire, such as those produced by low-severity prescribed fires and Indigenous cultural burns, may support microbial community resilience and maintain beneficial fungal symbionts capable of aiding seedling establishment. This has direct implications for forest management strategies that seek to balance fire risk mitigation with ecosystem recovery and resilience. Future work should expand on these findings through field-based experiments testing how microbial legacies, functional gene expression, and stress priming interact in varying environmental conditions within post-fire forest soil ecosystems. Understanding how to restore or harness beneficial soil microbial communities may be key to improving forest restoration outcomes in increasingly fire-prone landscapes.

## Abbreviations

ABA: Abscisic acid
EMF: Ectomycorrhizal fungi
Fv/Fm: Maximum quantum efficiency of photosystem II
ITS: Internal transcribed spacer
PSII: Photosystem II

